# Postural sway during single-legged standing is dependent on the body position assumption strategy and supports the stability and mobility trade-off hypothesis

**DOI:** 10.1101/215905

**Authors:** Bogdan Bacik, Grzegorz Sobota, Artur Fredyk, Kajetan J. Słomka, Grzegorz Juras

## Abstract

Studies that would address the problem of balance in the context of either the preceding or subsequent movement are scarce. We would like to propose the inverse relationship between stability and mobility as a general principle and that depending on the preceding action the postural sway characteristics in quiet standing would significantly differ. Thirteen female ballet school students were examined. Their mean age was: 15.1±0.95 years and the total of about 2500 (5^th^ school grade) and 4200 (6^th^ school grade) hours of dance training. The force plate was used to register ground reaction forces and moments. During the 30s trials subjects were to assume the *Passé* position in gait initiation (G-I) and gait termination (G-T) task condition. The following parameters were analyzed after restabilisation point estimation: the range of COP position, standard deviation of COP position, mean velocity in both planes and mean resultant velocity. The results showed significant differences between the G-I and G-T tasks in postural sway characteristics and confirmed our hypothesis. The practical implication of our study is the way of preparation to test position (PTTP) strategy might be as important as the measurement itself in the posture study protocols.

## 1. Introduction

The neural control mechanisms of human balance are still not fully explored despite the extensive studies in the area of biomechanics and related fields. A substantial progress in understanding balance has been made by identifying some general strategies and proposition of descriptive models of postural stability. Two terms tend to emerge in almost every posture study, which are stability and mobility. There is intrinsic trade-off between them and one cannot exist without the other. We would like to propose the inverse relationship between stability and mobility as a general principle. Although the stability/mobility continuum have been considered in different settings [1] we did not find postural control studies that would address the problem of balance in the context of either the preceding or subsequent movement. For example in sport competition an unstable position of a sprinter in starting blocks is conditioned by the imminent proceeding action which is a rapid start. In general, “starting positions” tend to maximise mobility while “final positions” optimise stability. As the role of mobility depends on the planned motor task, the question is whether it is possible that the same position has different effective postural control characteristics dependent on preceding action. While mechanical properties of the equilibrium remain the same, it conceivable that the control mechanism vary, which may affect the instantaneous ability to maintain balance. Based on the idea that the stability and mobility are inversely related [2] we would like to propose hypothesis that depending the preceding action the postural sway characteristics in quiet standing would significantly differ.

The magnitude of body sway provides a simple measure of balance and is commonly used in the clinical examination of postural control. The Romberg’s test, presented in the landmark study, is still one of a most common tools for detection of impairments in postural control in clinical environment. The interpretation of Romberg’s test results show that an increased postural sway is considered a sign of impaired postural control. Currently, more sophisticated methods are available (e.g. force platforms, EMG, Motion Capture Systems) and it is possible to show effects of different factors affecting postural sway in bipedal standing. Many factors have been studied and have different influence on postural sway. To count only few we have studies focused on feet position [3], light touch [4], visual information [5], motor imagery [6], age [7] and physical activity in daily life [8], [9]. Many of other factors and its influence on the human balance still remain unknown. The results concerning postural sway are not always straightforward to interpret, even within a homogeneous group [10]. Due to the abundance of factors influencing postural control, researchers carefully design experimental conditions and precisely define the body position. Nonetheless, the way the examined position is reached during examination is usually neglected. So far, not a single study has analysed the effects the way of assumption of the desired position prior to the examination.

The differences between the tasks conditions is found in the way the subject achieve examination position, this preceding action is called ‘preparing to test position (PTTP) strategy’. What is important, the final position is the same. The PTTP strategies are inherently linked to automated locomotion patterns, and we distinguish gait initiation (G-I) or gait termination (G-T) strategies (the tasks are explained in detail in Sec. 2.3). In order to realize experimental task of G-I and G-T we have decided to focus on one-legged stance. Even though they are both inherently linked to automated locomotion patterns, both of them are rather challenging for balance [11], especially compared to the bipedal quiet standing [12]. Gait initiation is difficult due to the transition from a quasi-static bipedal phase to a dynamic single-support phase. Gait termination requires slowing down the body to quiet standing. Additionally, during single-leg stance the base of support and the integrative information for interlimb coordination is decreased [13], therefore such position is difficult to maintain. The single leg stance has been described as a “quasi-static posture, as the body is in continuous motion and never achieves absolute equilibrium, even when the task is to remain as still as possible" [14]. The experiment requires one-leg standing, which is appropriate to select subjects who feel comfortable in the position and can maintain this posture for longer time. In consequence, we chose classical ballet dancers due to their high level of postural control and a great sense of balance [15] especially in the specific balance conditions of classic dance [13] [16]. The one-leg stance is close to the specific ballet position *Pass*é (Fig. 1). *Passé* appears in various ballet settings with different stability and mobility emphasis, e.g. in pirouettes a dancer needs to stabilize *Passé*. During *Passé Développé, the Passé* position is used as an intermediate position facilitating mobility. With this study we postulate that the expected outcomes of the preceding action on resultant postural sway characteristic will be different and will be dependent on this action. Therefore, the aim of this study is to explore the influence of body position assumption strategy on body postural sway.

**Figure 1.**
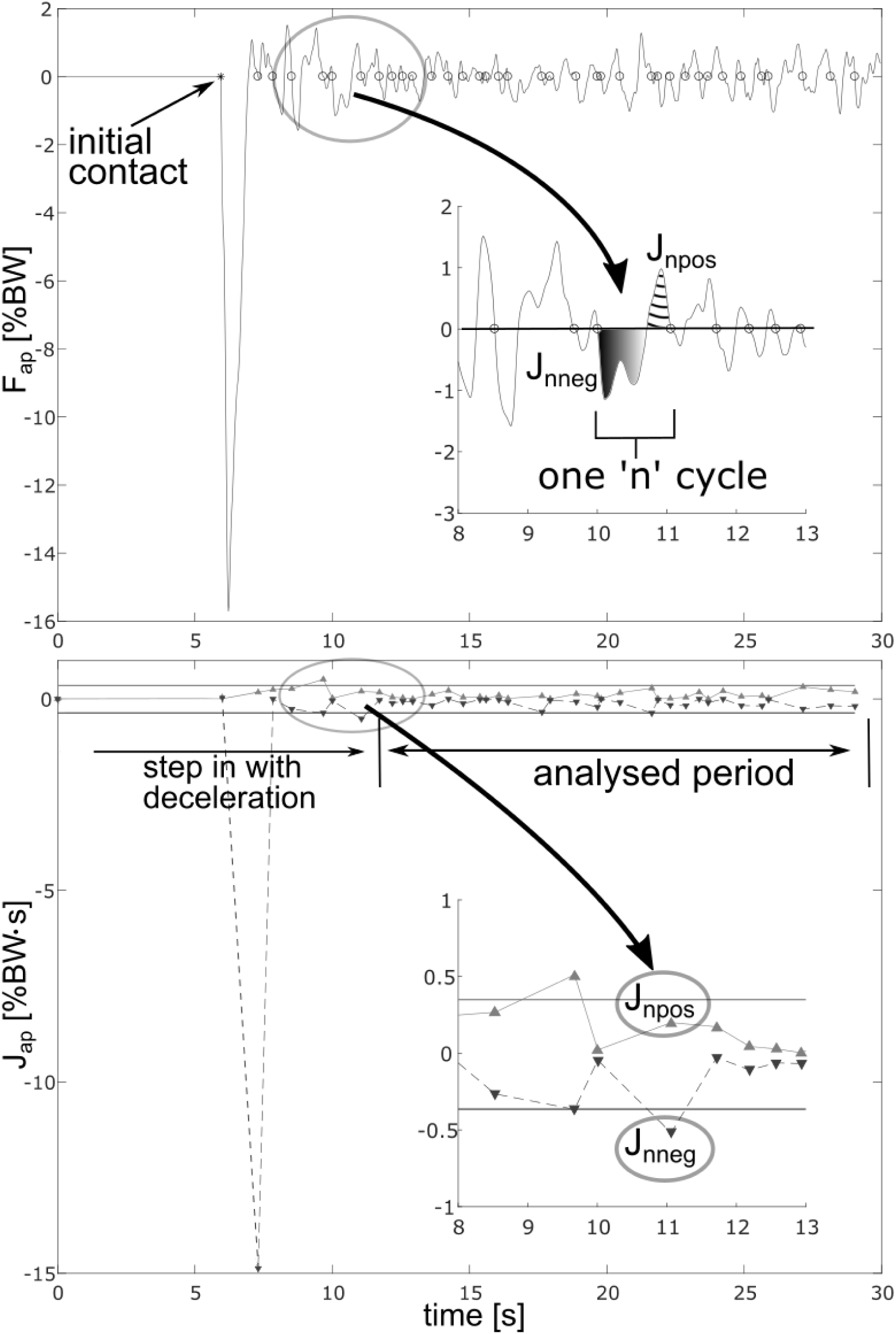
An example of force (top panel, anterio-posterior direction, F_ap_) and impulse(bottom panel, J_ap_) time series of one G-T trial (see Subsection 2.4.2). Zoomed parts correspond to positive and negative force and impulses for one of cycle, respectively. Note large negative impulse of body deceleration and the positive impulse of first correction after step in phase.

## 2. Methods

### 2.1. Subjects

Thirteen female students of 5^th^ (n=6) and 7^th^ (n=7) grade of the National Ballet School in Bytom (Poland) participated in the experiment. Professional training in this kind of school is intensive: 20 hours per week in 5^th^ grade and 22 hours per week in 7^th^ grade. At the time of examination, they had already undertaken the total of about 2500 and 4200 hours of dance training, respectively. Mean age, body mass and body height were respectively: 15.1±0.95 years, 50.1±5.42 kg and 163, 5±4,1 cm. The subject’s leg preference was examined with the Waterloo Footedness Questionnaire ([17], and all participants were right footed. None of the dancers reported any musculoskeletal disorders nor had an injury in the last three months before the examination. The study was approved by the Institutional Review Board and it was conducted in accordance with the Helsinki Declaration. Additionally, the approvals of parents and legal guardians as well as the school headmaster were obtained.

### 2.2. Instrumentation and set-up

To collected ground reaction forces and moments we have used a force platform (AMTI, Accugait) with 100Hz sampling frequency. The manufacturer’s software Netforce v. 3.5.3 was used in data collection. The surface of the platform was placed on the ground level to facilitate the execution of the experimental protocol.

### 2.3. Experimental Protocol

The measurement protocol consisted of four 30s trials. The subject’s task was to assume the *Passé* position (Fig. 1) in two different conditions, and maintain it till the end of the trial. All trials were done barefoot. Each of the subjects was required to achieve *Passé* position in gait initiation (G-I) and gait termination (G-T) task. The task conditions were examined separately on the right and left leg in random order. Each task on each leg was repeated twice for each subject giving in all a series of four trials. In the G-I task condition the one leg stance (*Passé* position) was reached as the end position starting from two legged quiet stance (i.e. required the lifting of a single leg). The G-T task conditions required from the participants to assume the *Passé* position (i.e. end position) after step forward from the position of two legged stance. All trials were done with eyes open. To avoid fatigue or boredom there was a 30s rest between trials [18]. The same measurements were repeated after 2, 3, 6 and 12 weeks. Not all dancers took part in all 5 measurement sessions, 1 missed 2 sessions and 6 of them missed one of the sessions. All measurements were carried out by the same investigator.

### 2.4. Data Processing and Analysis

#### 2.4.1. Preprocessing

A lowpass (10Hz) Buterworth filter (2^nd^ order) with no phase shift was applied to the signal. Using the measured ground reaction forces and force moments, the center of foot pressure (COP) was calculated with Matlab (Mathworks, USA).

#### 2.4.2. Restabilisation point estimation

In each direction (AP/ML), the force signal alternates between positive and negative values. Entire trial was divided into cycles defined as time intervals bounded by two consecutive zeros of the force curve (Fig. 2, top panel). Each cycle encompasses one negative period (negative values of force) and one positive (positive values of force). Positive (J_pos_) and negative (J_neg_) force impulses in each cycle were calculated by integrating over the cycle time (Fig. 2, top panel, zoomed part). A typical time course of force impulses is presented in Fig. 2 (bottom panel). Large negative force impulse after stepping on the force plate corresponds to deceleration of the body followed by a substantial first correction. Subsequently we observe mixed force impulses, what is typical due to body sway. We assumed that the step in motion was ended (restabilisation point) when the magnitude of force impulses (negative and positive) does not exceed the threshold for at least two seconds (Fig. 2, bottom panel, zoomed part). The upper and lower threshold levels were set as mean value of impulses from all the cycles within last five seconds of the trial increased by two standard deviations. A similar procedure was used by Singer [19], where they searched the “restabilisation point” using COP velocity. For G-T task the restabilisation point was calculated for anterio-posterior direction and for G-I task the restabilisation point was defined analogously using forces in the medio-lateral direction.

**Figure 2.**
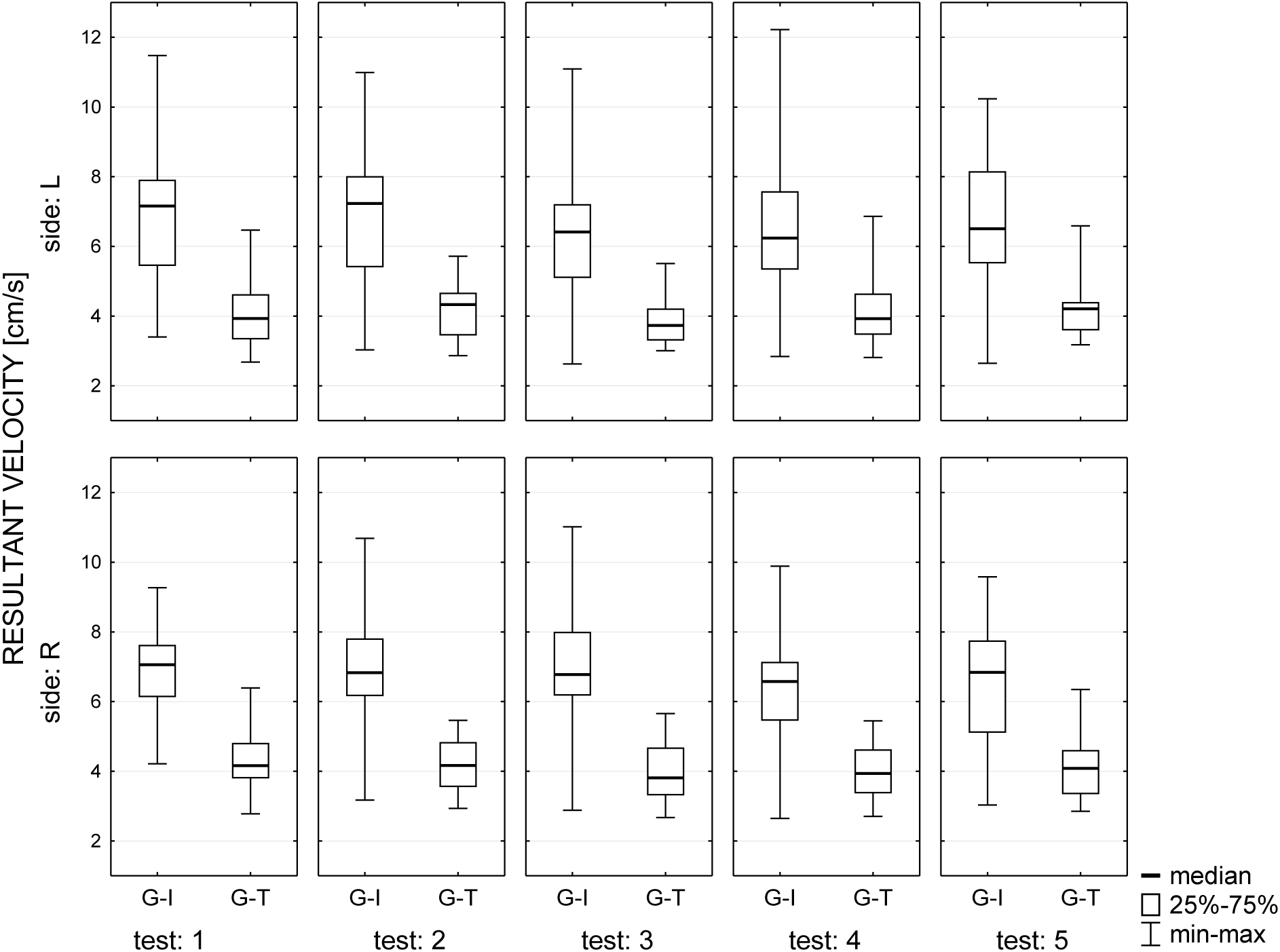
The differences in resultant velocity for both tasks for both sides (L/R) separately. Higher resultant velocity for G-I was observed throughout all 5 measurement sessions spread across 12 weeks of experiment.

#### 2.4.3. Body sway parameters

The body sway analysis was done for the time window from the restabilisation point to the end of a trail. The duration of analyzed time interval was similar for each subject, typically 20s (±1.39s). We have calculated the following parameters: the range of COP position, standard deviation of COP position, mean velocity in both planes and mean resultant velocity.

### 2.5. Statistical Analyses

The Shapiro–Wilk test and Levene’s test were performed to assess the equality of variances and check the data distribution for normality. Due to the five series of measurement the Mauchly’s test of sphericity also was done. The level of significance was set at *p≤0.01*. The initial analysis did not show normal data distribution nor the homogeneity of variance, that is why we have used non-parametric Wilcoxon, Mann-Whitney U test and Kruskal-Wallis ANOVA in order to infer statistically relevant features of the body sway quantities in the examined population. All statistical analyses were conducted with the use of Statistica 9.0 software package (StatSoft, Polska).

## 3. Results

### 3.1. Reliability of the testing procedures - ICC study

The intraclass correlation analysis ICC showed satisfactory reliability levels for all body sway parameters (Tab. 1). This proves that body sway parameters we proposed consistently quantify body sway characteristics of a subject. The lowest reliability levels (good, ranged 0.6 to 0.74) were observed for some of the amplitude parameters, while all the velocity parameters showed reliability on the excellent level (0.75 to 1.0). It is in accordance with the results of Pinsault and Vuillerme [20] and Salavati [21] who report that COP velocity is more reliable than all other measures.

**Table 1.** Intraclass Correlation Coefficients based on different sway parameters in both directions (AP, ML) and both PTTP strategies. High values of ICC imply good or excellent reliability.

|  | G-I |  | G-T |  |
| --- | --- | --- | --- | --- |
|  | *ICC A/P* | *ICC M/L* | *ICC A/P* | *ICC M/L* |
| COP range | 0,75 | 0,67 | 0,68 | 0,6 |
| Sd COP | 0,79 | 0,7 | 0,8 | 0,67 |
| COP velocity | 0,9 | 0,9 | 0,85 | 0,83 |
| Resultant velocity | 0,91 |  | 0,85 |  |

### 3.2. Differences between body sway for legs (right/left) and consecutive measurement sessions

The Mann-Whitney U test showed no significant effect of support leg at final position on body sway parameters. This allowed us to check the differences between PTTP strategies without differentiation between legs. No differences were observed between five consecutive measurement sessions (Kruskal-Wallis ANOVA, Fig. 3) for all variables. The observed lack of difference in postural sway between the dominant and non-dominant legs is in accordance with many studies reviewed by Clifford & Holder-Powell [12], conducted on healthy asymptomatic individuals.

**Figure 3.**
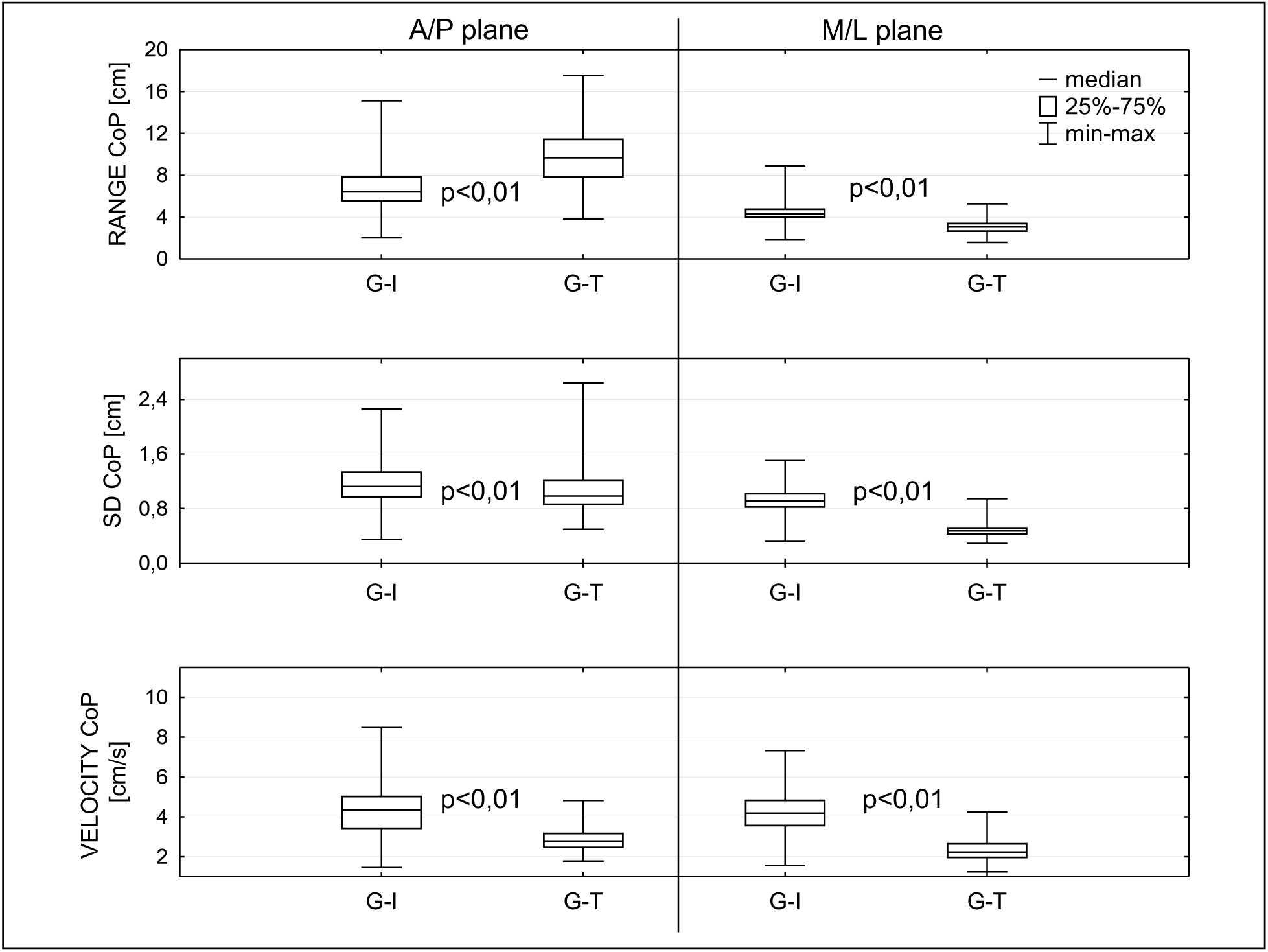
Range, SD and velocity in sagittal plane (AP) and frontal plane (ML) in gait initiation (G-I) and gait termination (G-T) tasks. In all cases the differences between PTTP strategies were significant (p< 0.01, Wilcoxon test).

### 3.3. Differences in body sway between specific types of PTTP strategy

The Wilcoxon test showed statistically significant differences between all analyzed body sway parameters (Fig. 4) for G-I and G-T strategies. All but one (the range of motion in the A/P plane which can be affected by a single stochastic event) were lower in magnitude in the G-T task. Standard deviation of COP displacement and its mean velocity are lower even in the A/P plane, which ascertains that the differences cannot be attributed to initial deceleration. In other words, it reassures that by introducing the notion of restabilisation, we successfully discarded mechanical effects of the initial step. Consequently, we propose interpreting the results in the context of *stability-mobility hypothesis* outlined in the introduction. *Passé* position is characterized by less body sway in the G-T task when the subjects aim to maximize stability of the new position. Conversely, increased body sway in the G-I scenario can be explained by the intention to increase mobility (leg raising is a part of the gait initiation motor program). Remarkably, the differences are maintained in the follow-up measurement sessions, and the values of the parameters are stable (Fig. 3). Figure 4 depicts lower SD and mean velocity for G-T even in the AP plane, which means that the effects of initial deceleration were successfully removed.

## 4. Discussion

The main aim of the present study was to test whether the way we achieve an examination position would lead to changes in postural sway. The results showed significant differences between the G-I and G-T tasks and confirmed our hypothesis. The body sway velocity in the AP and ML direction are of the same order of magnitude as those reported for one-leg stance of young healthy subjects [18]. It should not be surprising that subtle factors affect body sway just like in Fawver’s et. al. [22] study. They showed that movement of CoP can be increased by presenting emotional images to the subjects. The repeatability of our results obtained in each of the five remote testing sessions (Fig. 4) is to confirm the existence of the automatic mechanism associated with the sequence of starting or stopping the motion of ballet dancers. We propose to attribute this effect to the acclaimed principle of *stability-mobility conservation* in balance, however further studies would be required to probe generality of that characteristic across different study groups and postures.

There is still no consensus about how the ballet training affects balance abilities, so it is possible that our results heavily depend on the choice of study group. On the one hand, some studies indicate that dancers are characterized by less body sway [16] on the other hand there are reports suggesting higher values of COP parameters [23]. Indeed, dance training may reduce the number of constraints on ankle–hip coordination in order to enhance adaptability and flexibility of dance-specific movement patterns [24]. Moreover, it has been suggested that the ballet dancers use different coordination strategy [25] and some studies postulate that it mostly relies on visual afferences [23]. In addition, the dancers during their routine are used to finding their “centre” or “vertical” [13] which may increase the sensitivity of this group to the proposed test conditions. On the other hand, no differences in the parameters of body sway between healthy dancers and the control or players of different sports disciplines when standing with one leg were reported by some authors ([26] [23] [15]. Nonetheless, no much comparative studies between the dancers and other athletes have been conducted in the setting of unipedal stance with eyes open ([8].

The most practical implication of our study is the postulate of paying attention to the preparing to test position (PTTP) strategy in the stabilographic study protocols. This concerns especially single leg stance. One may speculate that careful prescription method of adoption of the examination position could improve the quality of the collected data.

**Picture 1.**
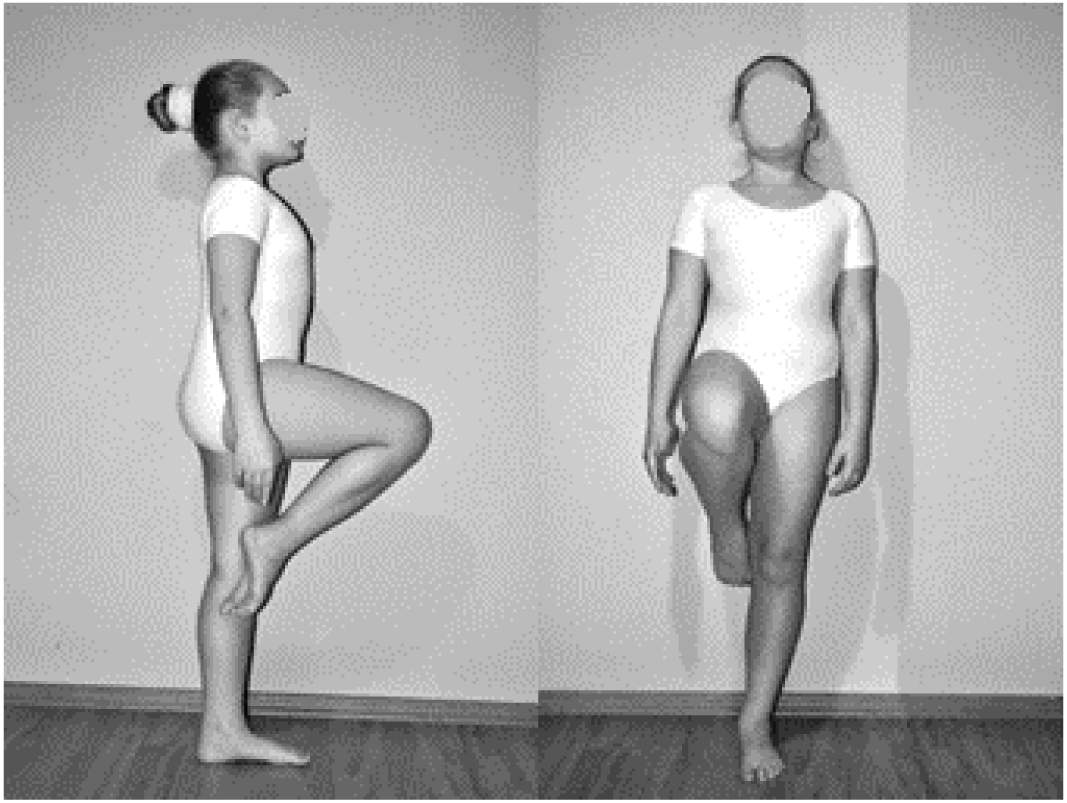
*Passé* position in classical ballet used in the experimental protocol; side and frontal view.

## Acknowledgments

The project was supported by Statutory Funds of Faculty of Physical Education, The Jerzy Kukuczka Academy of Physical Education in Katowice. We would like to thank Jim Usherwood and Slobodan Jaric for their valuable comments and helpful discussion.

## Conflict of interest statement

The authors have no conflicts of interest to disclose.

